# Transiently restricting individual amino acids protects *D. melanogaster* against multiple stressors

**DOI:** 10.1101/2024.04.04.588160

**Authors:** Tahlia L. Fulton, Joshua J. Johnstone, Jing J. Tan, Krithika Balagopal, Amy Dedman, Travis K. Johnson, Andrea Y. Chan, Christen K. Mirth, Matthew D. W. Piper

## Abstract

Nutrition and resilience are linked, though it is not yet clear how diet confers stress resistance or the breadth of stressors that it can protect against. We have previously shown that transiently restricting an essential amino acid can extend lifespan and also protect against nicotine exposure in *Drosophila melanogaster*, raising the possibility that amino acid restriction is geroprotective because of elevated detoxification capacity. Here, we sought to characterise the nature of this dietary mediated protection, and determine whether it was sex, amino acid, and/or nicotine specific. When we compared between sexes, we found that isoleucine deprivation increases female, but not male, nicotine resistance. Surprisingly, we found that this protection afforded to females was not replicated by dietary protein restriction and was instead specific to individual amino acid restriction. Other studies have documented methionine or leucine restriction conferring stress resistance, though we previously found that individually depriving them did not increase nicotine resistance. We therefore wondered whether reducing the severity of restriction of these amino acids could confer nicotine resistance. This was true for methionine restriction, and we found that flies fed a diet containing 25% methionine for 7 days protected against subsequent nicotine poisoning (∼30% longer lived than controls with all amino acids). However, when dietary leucine was altered, nicotine resistance changed, but no single diet was protective. To understand whether these beneficial effects of diet were specific to nicotine or were generalisable across stressors, we pre-treated with amino acid restriction diets and exposed them to other types of stress. We did not find any diets that protected against heat stress or infection with the bacterium *Enterococcus faecalis*. However, we found that some of the diets that protected against nicotine also protected against oxidative and starvation stress, and improved survival following cold shock. Interestingly, we found that a diet lacking isoleucine was the only diet to protect against all these stressors. These data point to isoleucine as a critical determinant of robustness in the face of environmental challenges.

## Introduction

Nutrition is closely linked to health and the more we explore this connection, the closer we come to being able to deliberately manipulate components of the diet to influence health and fitness. Nutrient restriction, in particular protein restriction, has been linked to increased lifespan, improved metabolic health, and stress resistance in many model systems, usually at the cost of reproductive output (Zhang et al. 2023; Krittika and Yadav 2020; Jongbloed et al. 2017; Robertson et al. 2015; Mirzaei, Suarez, and Longo 2014; Brandhorst et al. 2013; Soultoukis and Partridge 2016). These effects can also be mimicked by restricting dietary amino acids (Yap et al. 2020; Yeh et al. 2023; Trautman, Richardson, and Lamming 2022; Grandison, Piper, and Partridge 2009; Fontana et al. 2016). For instance, transient isoleucine deprivation enhances nicotine resistance and extends lifespan in *Drosophila* (Fulton, Mirth, and Piper 2022), methionine restriction enhances resilience to chemical and thermal stress in yeast, mice, and human cells (Trocha et al. 2019; Johnson and Johnson 2014; Miller et al. 2005), and depriving mice of tryptophan protects them against ischaemic reperfusion injury (Peng et al. 2012). These beneficial effects of diet, particularly these short-term restrictions, have attracted interest for their potential to enhance human health. Current data indicate that evolutionarily conserved mechanisms mediate these effects (Harputlugil et al. 2014; Peng et al. 2012; Fulton, Mirth, and Piper 2022), though little is known about the breadth of stress resistance these dietary manipulations afford and/or if there are costs beyond reproductive arrest that are associated with their implementation. These considerations, as well as understanding their mechanisms, are important when seeking to apply treatment protocols across species.

It is important to consider that seemingly beneficial dietary strategies may impose unintended costs, or trade-offs. One trade-off that is well described in the literature is between lifespan and reproduction; while lifespan is longest on low protein, high carbohydrate diets, reproduction in females is highest in intermediate protein, intermediate carbohydrate diets (Lee et al. 2008; Holliday 1989; Partridge, Gems, and Withers 2005). Because these traits are optimised on different diets, this makes it difficult for animals to be both long lived and have maximal reproduction. Using a similar logic, if short-term amino acid restriction can protect flies against nicotine, and prolong life, does this mean that it will protect them against other stressors too, or do these benefits trade-off against other dimensions of stress resistance?

There are different types of biotic and abiotic stressors against which organisms must evolve strategies to resist, tolerate and/or avoid. In nature, flies and other animals frequently encounter stressors such as naturally occurring insecticides, temperature fluctuations, and infections (Kaunisto, Ferguson, and Sinclair 2016). It is also likely that an animal will encounter more than one stressor at a time. For example, winter is not only cold but also dry, so insects must tolerate both simultaneously to survive (Sinclair et al. 2013). It is therefore important that organisms launch multiple resistance phenotypes, even when sensing only one stressor, so they have the best chance to survive current and anticipated environments.

In this paper, we present our findings on the way *Drosophila melanogaster* initiate a broad range of stress responses when experiencing short term amino acid deprivation. Our study contributes to a deeper understanding of the complex interactions between nutrition and stress resistance, offering insights that could inform the development of personalised dietary strategies for promoting health and longevity.

## Methods

### Fly husbandry

Experiments were conducted using white-eyed *Drosophila melanogaster* (strain Dahomey; wDah). Outbred wDah stocks are maintained in high-density population cages with continuous overlapping generations on a sugar yeast (SY) diet (Bass et al. 2007) (Supplementary table 1) at 25°C, ambient humidity, and a 12:12 hour light:dark cycle. Experimental flies were reared from egg to adult at a density of ∼250-300 flies per bottle with 70mL SY medium and mating status was standardised by keeping newly emerged adult flies in mixed cohorts on fresh SY diet for 2 days following eclosion (Figure 1) Unless specified, experimental flies were mated, female, wDah. Two days after eclosing, flies were lightly anaesthetised with CO_2_ and sorted by sex into vials containing a complete synthetic diet (Piper et al. 2014) (Supplementary table 2) in cohorts of 5 flies per vial. Flies were transferred to fresh food every Monday, Wednesday, and Friday, unless otherwise specified. Experiments and stocks were maintained at 25°C, 60% humidity and a 12:12 hour light:dark cycle.

**Figure 1:**
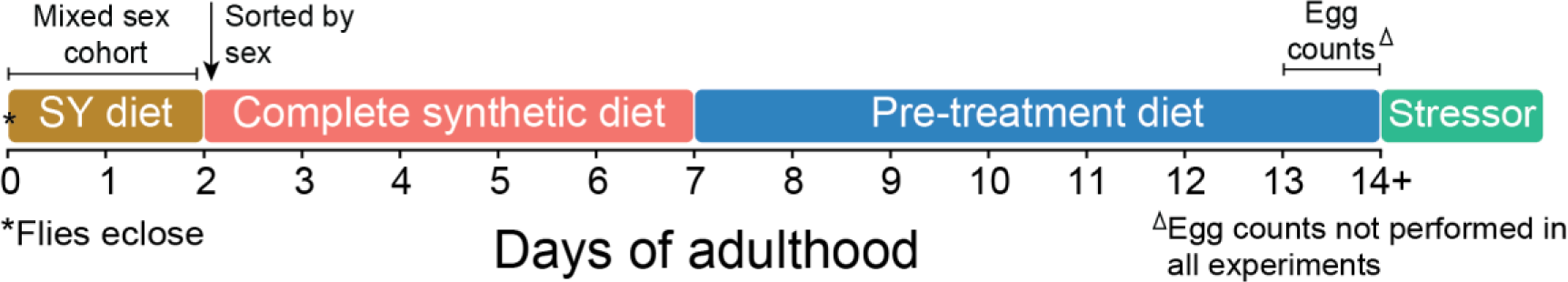
General methods for experiments. Newly emerged adult flies were transferred to fresh Sugar Yeast (SY) food to mate for 2 days, then sorted into vials containing a complete synthetic medium at a density of 5 flies (of the same sex) per vial. Unless otherwise specified, flies were transferred to their pre-treatment diet on day 7 of adulthood. If fecundity was measured before exposure to the stressor, flies were transferred to fresh vials on day 13 of adulthood and the eggs in those vials were counted on day 14.

### Synthetic media preparation

Synthetic media were prepared as described by Piper et al. (2014) containing the nutrients listed in Supplementary table 2. The complete synthetic diet contained an exome matched ratio of amino acids (Piper et al. 2017) (FLYaa), and the other pre-treatment diets were prepared in the same way except with a reduced amount of the focal amino acid. Media were prepared in advance and stored for up to 4 weeks at 4°C.

### Laced-media preparation

Survival under nicotine or paraquat exposure was measured using fly food that was laced with the respective drug. Nicotine-laced medium was prepared by aliquoting 100µL of diluted free base nicotine (Supplementary table 3) into a vial containing 3mL of cooled, gelled complete synthetic medium (final concentration of 0.83mg/mL nicotine in vials). Paraquat laced vials were similarly prepared by aliquoting 100µL of diluted methyl viologen dichloride hydrate (Supplementary table 3) onto 2mL of cooled, complete synthetic medium (final concentration of 10mM paraquat in vials). Laced vials were then kept in a fabric cover at room temperature for 24-48h to ensure an even dispersion of toxin throughout the food. Laced food was prepared only in sufficient volumes to match what was needed for immediate use, and only 24-48h in advance of use, as these drugs lose potency over time.

### Egg counting

When measuring fecundity, flies were first allowed to lay eggs in fresh vials for 24h. Following this, flies were transferred to new vials and the vials containing eggs were photographed using a stereo microscope (Zeiss; Stemi 508) with an attached camera (Zeiss; Axioxam ERc 5s). The photographs were then piped through an application that was made in-house by Jing J. Tan to count the number of eggs in each vial. This application is available for public use here: https://github.com/Eyehavelived/egg_counter

### Nicotine and paraquat exposure protocol

Following pre-treatment (Figure 1), cohorts of 5 flies per vial were transferred into vials containing either 0.83mg/mL nicotine or 10mM paraquat. After 48h in these toxin-laced vials, flies were transferred to freshly prepared toxin-laced vials. Once initially exposed to their respective toxin, survival was recorded at 7am, 1pm and 7pm for 3 days, and then at 8am and 5pm for 2 days, following which any remaining surviving flies were censored. Fly survival was recorded using the software DLife (Linford et al. 2013).

### Starvation protocol

After they were pre-treated with amino acid deprivation (Figure 1), flies were mildly anaesthetised with CO_2_ (3L/min for < 5 min) and placed individually into wells of a 96 well tissue culture plate (Falcon: FAL353072) that contained 700uL of 2% agar (Sigma Aldrich: A7002). Before flies awoke, the lid was placed on the plate to contain single flies to individual wells. We fitted a custom-built plate scanning robot with a digital camera (Dino-Lite) and used this robot to photograph each well of each plate every hour. We sorted the images by well location and used the photos in series to determine when the fly stopped moving, at which point they were scored as dead.

### Heat knockdown protocol

Following pre-treatment with their respective diets (Figure 1) flies were individually transferred to 5mL glass vials using a mouth-controlled aspirator. The vials were then submerged into a 39°C recirculating water bath that was heated by a digital thermos-regulator (Model: TH5; Ratek). Heat knockdown time was measured as the time taken, to the nearest second, until a fly was immobile.

### Cold shock protocol

Flies were pre-treated with their respective diets (Figure 1) and then individually transferred to 1.5mL Eppendorf tubes using a mouth-controlled aspirator. The Eppendorf tubes were then submerged in a tank containing 50% propylene glycol (v/v, with water) that had been cooled to 0°C using a Thermoline liquid cooler (TRC-500). After 5h at 0°C, flies were then transferred to 25°C to recover. Recovery time was measured as the time taken (to the nearest second) for flies to begin moving again. A small number of flies (11 out of 300) did not recover within 2 hours, and they were considered dead and omitted from the analysis. All flies that recovered were transferred individually into vials containing a complete synthetic diet, and survival was recorded 2-3 times a day for 5 days using the software DLife (Linford et al. 2013).

### Infection protocol

Wild-caught *E. faecalis* stocks (Lazzaro, Sackton, and Clark 2006) were stored at - 80C in Luria Bertani (LB) broth with 15% glycerol. To prepare bacteria to infect flies, a stock was first streaked onto an LB plate and grown overnight at 37C. A single bacterial colony was then picked from this plate and grown overnight in 2mL of LB broth, in an orbital shaker kept at 37C rotating 200 times per minute. The overnight cultures were diluted with sterile phosphate buffered saline (PBS, Ph = 7.4) to an optical density (OD_A600_) of 0.1. Pre-treated flies (Figure 1), were lightly anaesthetised with CO_2_ (3L/min for <5min) and injected with 0.1 OD_A600_ *E. faecalis* using the septic pinprick method (Khalil et al. 2015). Control flies were instead injected with sterile PBS. Following infection, flies were transferred in cohorts of 5 to a complete synthetic diet, and survival was recorded 2-3 times a day using the software DLife (Linford et al. 2013). Infected and control flies were transferred to fresh food every day, and the old vials were photographed to measure daily fecundity.

### Statistical analysis

All analyses were completed using R (Team 2021) (version 4.2.2) and R Studio (Team 2020) (version 1.4.1106) and we created all plots using ggplot2 (Wickham 2016). All data and scripts are publicly available at: [To be made freely available through Figshare on publication].

To determine whether the independent variables of a model could explain variation in the data, we initially analysed the models using a type II or III ANOVA from the package car (Fox and Weisberg 2019).

Cox Proportional-Hazards modelling was used to analyse survival. To do this, we used the “coxph” function from the package survival (Therneau 2021).

Differences in fecundity were analysed using a linear model (base R, “lm”) and post-hoc comparisons from the emmeans (Lenth 2021) and multcomp (Hothorn, Bretz, and Westfall 2008) packages.

Linear models (base R, “lm”) were also used to model both duration of pre-treatment and dose of amino acid as a function of survival. To determine whether pre-treatment duration or availability of the focal amino acid significantly impacted survival, we analysed the model using the “Anova” function from the package car (Fox and Weisberg 2019).

## Results

### Pre-treatment diets that protect female flies from nicotine do not protect males

In our previous work, we found that some diets lacking an essential amino acid protected female flies from subsequent nicotine poisoning (Fulton, Mirth, and Piper 2022). While female flies stop laying eggs when they are fed food without one or more essential amino acids, this protective effect cannot be attributed solely to a simple trade-off against reproduction. This is because neither a leucine or methionine dropout, or a diet missing all amino acids was protective, yet they still reduced egg laying to the same level as protective diets (Fulton, Mirth, and Piper 2022). As protection by diet cannot be simply explained by a trade-off against reproduction, we hypothesised that males – who are thought to expend less energy for reproduction (Magwere, Chapman, and Partridge 2004) - could be afforded the same protective benefits of diets that increased female flies’ nicotine resistance. To explore this, we pre-treated both sexes with a diet lacking isoleucine for 7 days, which we previously found to be the most protective pre-treatment regimen (Fulton, Mirth, and Piper 2022). We also pre-treated flies with a nutritionally complete diet to normalise results between males and females, and a protein-free diet to control for removing all amino acids.

When comparing males and females that were not pre-treated (complete diet), males had a greater nicotine resistance than fully-fed females (Figure 2. A; P = 0.04). However, pretreating flies with an isoleucine dropout diet for 7 days increased female, but not male, nicotine resistance (Figure 2. B; Table 1). Removing all amino acids was equally detrimental to both sexes (Table 1). These results indicate that there is sexual dimorphism in the stress resistance afforded by diet.

**Figure 2.**
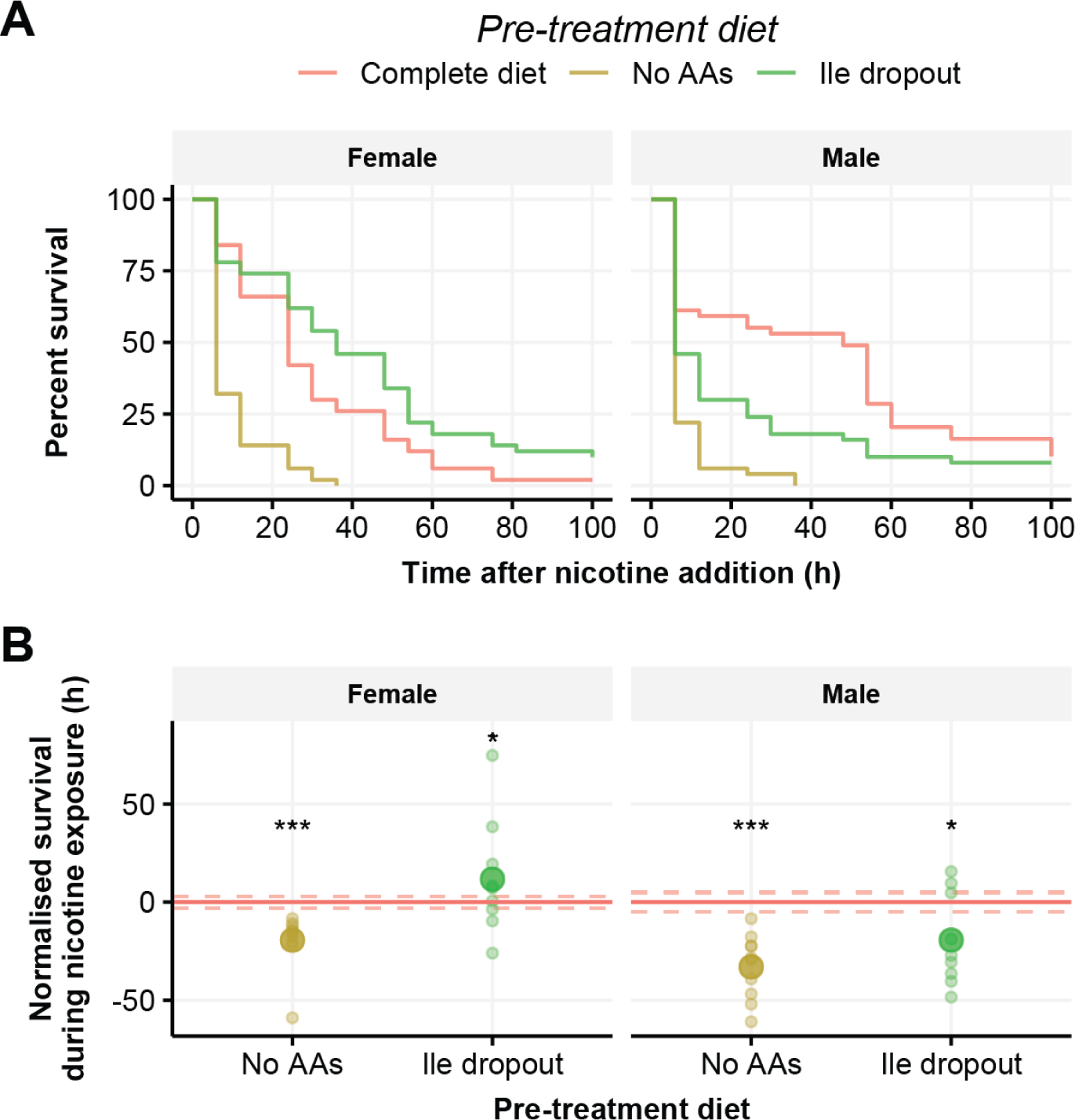
Pre-treatment with a diet lacking isoleucine protects female, but not male, flies against nicotine poisoning. Male and female flies were pre-treated with one of three diets - isoleucine dropout, no amino acids (AAs) or a complete diet - before chronic exposure to 0.83mg/mL nicotine. (**A**) Survival curves of flies immediately after introduction of nicotine. (**B**) Survival time of pre-treated flies that has been normalised to survival of flies fed a nutritionally complete synthetic diet (red horizontal line +/− SE indicated by dashed red lines), small circles represent mean lifespan for each replicate and large circles representing the group mean. In females, flies that were pre-treated with a diet lacking amino acids had reduced nicotine resistance (P < 0.001), whereas pre-treatment with an isoleucine dropout diet protected flies from nicotine (P = 0.02). Compared to the control diet, male flies were more susceptible to nicotine when they were pre-treated with either a diet lacking amino acids (P < 0.001) or isoleucine alone (P = 0.02). N = 49-50 flies per pre-treatment group for each sex. ***P < 0.001, ** P < 0.01, *P < 0.05.

**Table 1.** Differences in survival between flies that were pre-treated with a diet lacking amino acids (no AAs) or an isoleucine dropout diet compared to flies that were fed a complete diet, separated by sex. Summary of cox-proportional hazards modelling. Confidence level = 95%.

| Terms | n | Estimate | Z value |
| --- | --- | --- | --- |
| **Female** |  |  |  |
| Complete diet | 50 | NA | NA |
| No AAs | 50 | 1.339 | **6.03\*\*\*** |
| Ile dropout | 50 | -0.477 | **-2.27\*** |
| **Male** |  |  |  |
| Complete diet | 49 | NA | NA |
| No AAs | 50 | 1.215 | **5.24\*\*\*** |
| Ile dropout | 50 | 0.483 | **2.26\*** |
| ***p < 0.001, **p < 0.01, *p < 0.05 |  |  |  |

### Restricting individual amino acids, but not all amino acids proportionately, is protective against nicotine exposure

Restricting protein in the diet of model organisms has repeatedly increased the consumer’s resilience to stress (Mirzaei, Raynes, and Longo 2016). We were therefore interested in whether protein restriction could protect female flies against nicotine in the same way that single amino acid deprivation does. In all prior experiments, we have found that completely depriving flies of protein for short durations has not been beneficial. However, we have also found that the strength of the protective response varies with the identity of the amino acid, the degree of restriction and the duration of pre-treatment (Fulton, Mirth, and Piper 2022). It is therefore possible that manipulating total protein could be protective when it is restricted in a way that we have not yet explored. We hypothesised that feeding flies less protein, or starving them of protein for a shorter period of time, could protect them against nicotine in the same way that restricting a single amino acid does.

To do this, we pre-treated two separate cohorts of flies. The first cohort was fed one of seven diets in which all amino acids were restricted to 75%, 50%, 25% or 0% of the amount in the complete diet for seven days prior to exposure to nicotine. The second cohort was fed a diet lacking amino acids/protein for shorter lengths of time before nicotine exposure. We found that both methods of protein restriction significantly impacted fecundity, which we measured in the 24h prior to nicotine exposure (Figure 1). We found that for every 6.66% that protein was reduced in the diet, egg laying was reduced by approximately 1 egg per female in the 24 hours measured (F_4_ = 81.49, P < 0.001). Similarly, consuming less protein by spending more time on a diet without protein also reduced fecundity by approximately 2 eggs per female over 24h for every additional day without protein (F_4_ = 76.79, P < 0.001). These decreases in fecundity are similar to what is observed when flies are deprived of a single essential amino acid (Alves et al. 2022). When we exposed these cohorts of flies to nicotine after pretreatment, we found that, contrary to expectations, no level of protein restriction increased nicotine resistance, and reducing protein to 25 or 0% was detrimental (Figure 3.C; Table 2; Supplementary Figure 1.A). Similarly, reducing the time that flies were starved of protein did not increase their nicotine resistance, and, as we previously found, starving them of protein for 7 days was detrimental (Figure 3.D; Table 2; Supplementary Figure 1.B). These results suggest that the benefits of individual amino acid restrictions prior to nicotine poisoning are specific to individual amino acids and perhaps require the presence of one or more of the remaining 19 amino acids.

**Figure 3.**
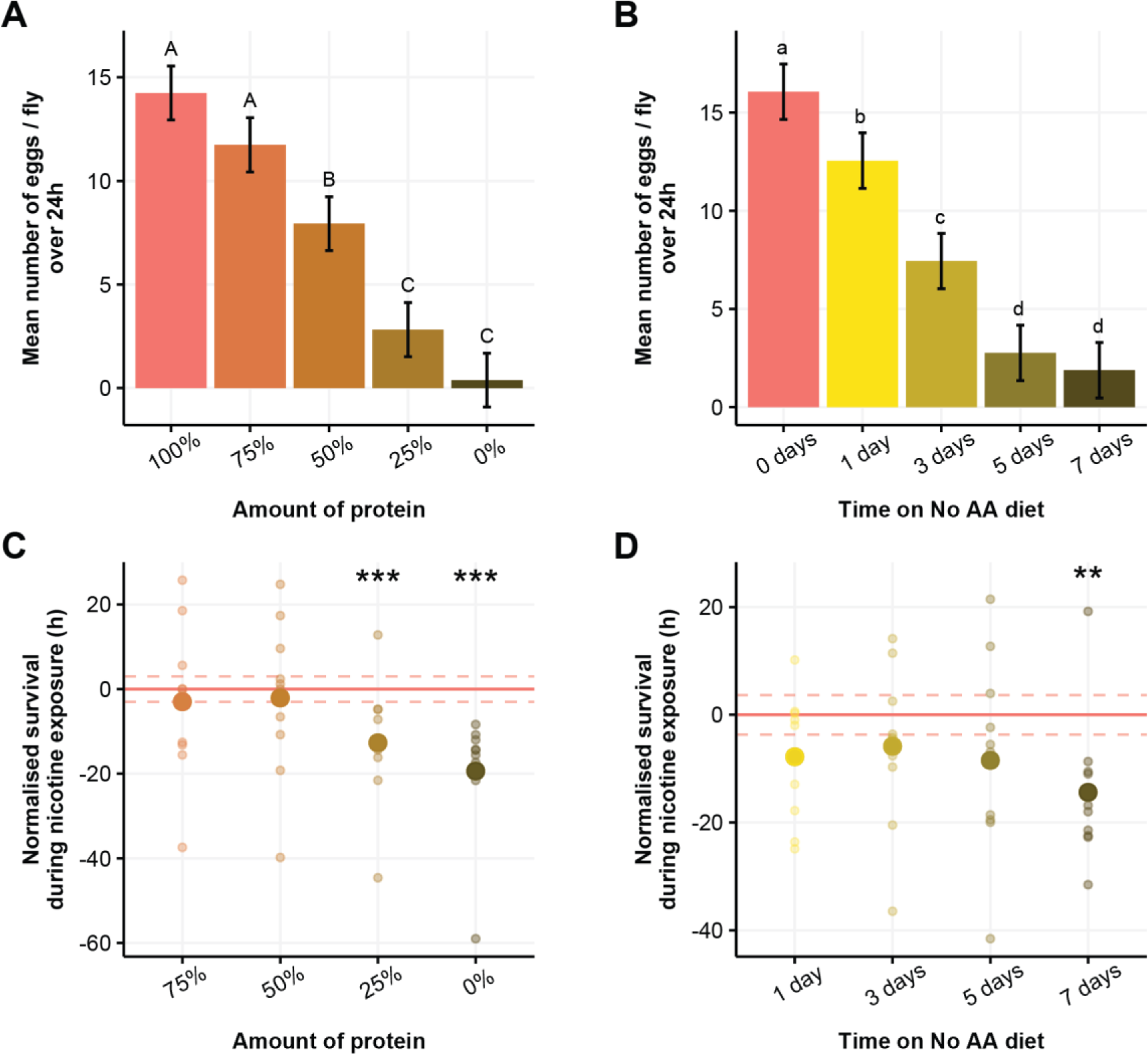
Restricting all amino acids does not protect flies against nicotine. Flies were either pre-treated with diets where all amino acids were reduced for 7 days, or where all amino acids were absent for 1, 3, 5 or 7 days before chronic exposure to 0.83mg/mL nicotine. (**A**) Mean number of eggs (+/− SE) per fly in the 24h before nicotine exposure when the concentration of amino acids is altered and (**B**) when all amino acids are removed for various days. (**C**) Survival time of pre-treated flies that has been normalised to survival of flies fed 100% of amino acids (red horizontal line +/− SE indicated by dashed red lines), small circles represent mean lifespan for each replicate and large circles representing the group mean. Flies tolerated nicotine less when they were pre-treated with diets that had 25% or 0% of the amino acids found in the complete diet (P < 0.001). (**D**) Survival time of pre-treated flies that has been normalised to survival of flies that were fed a complete diet (red horizontal line +/− SE indicated by dashed red lines), small circles represent mean lifespan for each replicate and large circles representing the group mean. Flies were less resistant to nicotine when they were fed a diet lacking all amino acids for 7 days before exposure. ***P < 0.001, ** P < 0.01, *P < 0.05.

**Table 2.** Differences in survival between flies that were pre-treated with diets that had reduced, or no protein compared to flies that were fed a nutritionally complete diet. Summary of cox-proportional hazards modelling. Confidence level = 95%.

| Terms | n | Estimate | Z value |
| --- | --- | --- | --- |
| **Protein restriction** |  |  |  |
| 100% protein | 50 | NA | NA |
| 75% protein (7 days) | 50 | 0.098 | 0.48 |
| 50% protein (7 days) | 50 | 0.102 | 0.5 |
| 25% protein (7 days) | 50 | 0.681 | **3.33\*\*\*** |
| 0% protein (7 days) | 50 | 1.273 | **6.03\*\*\*** |
| **No protein** |  |  |  |
| 100% protein | 50 | NA | NA |
| 1 day (0% protein) | 50 | 0.309 | 1.5 |
| 3 days (0% protein) | 50 | 0.296 | 1.44 |
| 5 days (0% protein) | 45 | 0.380 | 1.79 |
| 7 days (0% protein) | 50 | 0.633 | **3.05\*\*** |
| ***p < 0.001, **p < 0.01, *p < 0.05 |  |  |  |

### Assessing how availability and duration of methionine and leucine dilution impacts nicotine resistance

Given that restricting all amino acids proportionately did not protect flies from subsequent nicotine poisoning, we decided to return to exploring the benefits of restricting individual amino acids. In particular, we were interested in discovering the conditions that conferred the greatest benefit when restricting different amino acids. We previously found that the conditions that provided the maximum protection when isoleucine was altered were different to the conditions when threonine is altered (Fulton, Mirth, and Piper 2022). Specifically, removing isoleucine from the diet for 7 days was the most protective, whereas threonine only needed to be reduced to 25% to show the greatest protection against nicotine. In our initial screen, where we removed each amino acid individually, we found that there was no effect of depriving flies of either methionine or leucine, whereas removing any one of the other 8 essential amino acids for 7 days provided some degree of protection. This was particularly curious because other research describes the benefits from restricting these amino acids. Methionine restriction is strongly associated with longevity and increased metabolic health in flies, worms, and mice (Ables and Johnson 2017) and leucine is one of the three branched chain amino acids, and the other two, isoleucine and valine, both protected flies against nicotine when removed from the diet (Fulton, Mirth, and Piper 2022). As we have already shown that intensity of restriction and length of pre-treatment can impact the benefits of single amino acid restriction, we wanted to know whether there were conditions where flies were protected when we modified the availability of either methionine or leucine.

We individually restricted methionine or leucine to 75%, 50%, 25% or 0% of the amount in the complete diet. As we restricted the amount of these amino acids, we simultaneously modified duration of dietary pre-treatment to 7, 5, 3 or 1 day prior to nicotine exposure, resulting in a combination of 16 pre-treatment conditions plus a complete diet control. In the 24h immediately before nicotine exposure, we measured the fecundity of these flies to further understand the relationship between toxin resistance and reproductive output. When we modified methionine, we found that the only condition that significantly altered egg laying was a diet completely lacking methionine for 7 days (Figure 4. A; Table 3). Although methionine is an essential amino acid, this result was unsurprising because egg laying has previously been shown to exhibit a slower decline after methionine removal than that of the other amino acids (Alves et al. 2022). When we removed leucine from the diet, we observed a proportional reduction in egg laying, where fecundity was reduced by approximately 2 eggs per female in the 24h measured for every additional day that flies spent on a leucine dropout (Figure 4. B; Table 3; Supplementary table 4). Interestingly, we saw that a 25% leucine diet had a less steep decline in egg laying of approximately 1 egg/female/24h for every additional day spent on this diet and this decline appeared to level out after 5 days spent on the diet. Neither 50% (P = 0.14) nor 75% (P = 1) had a noticeable change in egg laying over time. Together, these data indicate that the flies were experiencing restriction for each of these amino acids to differing extents, and reinforces that they are likely to have internal reserves of amino acids on which they can draw to sustain reproduction (Johnstone et al. 2024).

**Figure 4.**
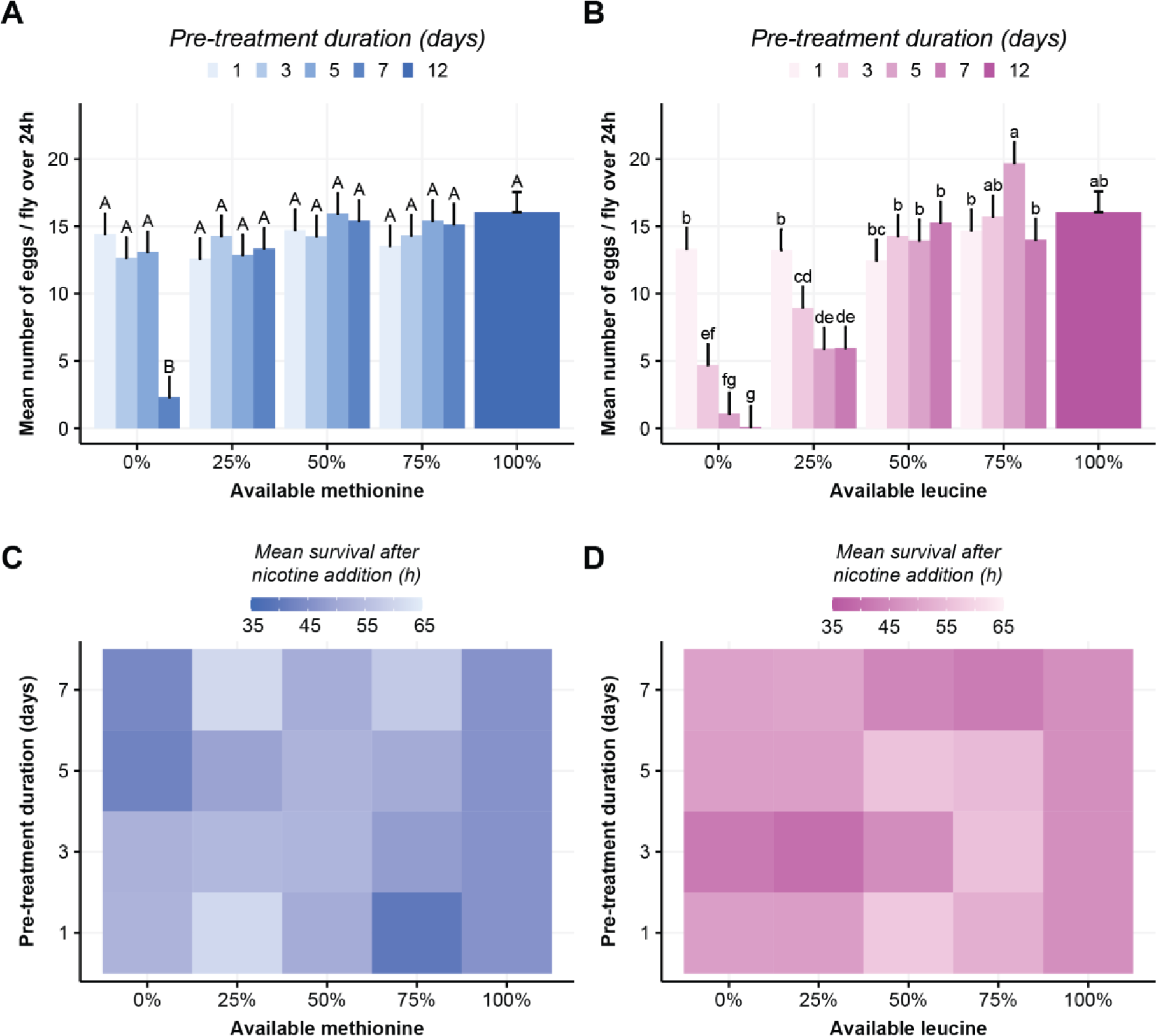
Amino acid identity and availability modify protection against nicotine stress. Flies were pre-treated with diets containing 1 of 5 amounts of either methionine or leucine (0%, 25%, 50%, 75%, 100%) for either 7, 5, 3, or 1 day before fecundity was measured and flies were chronically exposed to 0.83mg/mL nicotine. (A) When dietary methionine was modified, only a diet lacking methionine for 7 days reduced fecundity (B) However, restricting dietary leucine resulted in females laying approximately 2 fewer eggs/24h for every day she spent on a leucine dropout (95% CI [−2.57, −1.76]), 1 fewer egg/24h for every day she spent on a 25% leucine diet (95% CI [−1.64, −0.84]), or no decline over time when fed 50% or more leucine (P > 0.1). (C) Survival when methionine was modified was influenced by both the amount of methionine and the duration of pre-treatment (Table 4), with a 25% methionine for 7 days offering the greatest protection (P = 0.012) (D) Leucine dose (P = 0.02) but not pre-treatment duration (P = 0.59) significantly impacted survival on nicotine, although the longest lived flies - that were fed a 50% leucine diet for 5 days - were not protected (P = 0.09). N = 50 flies for each combination of pre-treatment duration and available amino acid. Pre-treatment with 100% of each amino acid could not differ in duration time and N = 50 flies for this group in total. The survival data has been normalised and represented in Supplementary Figure 2 to show variation in survival.

**Table 3.**
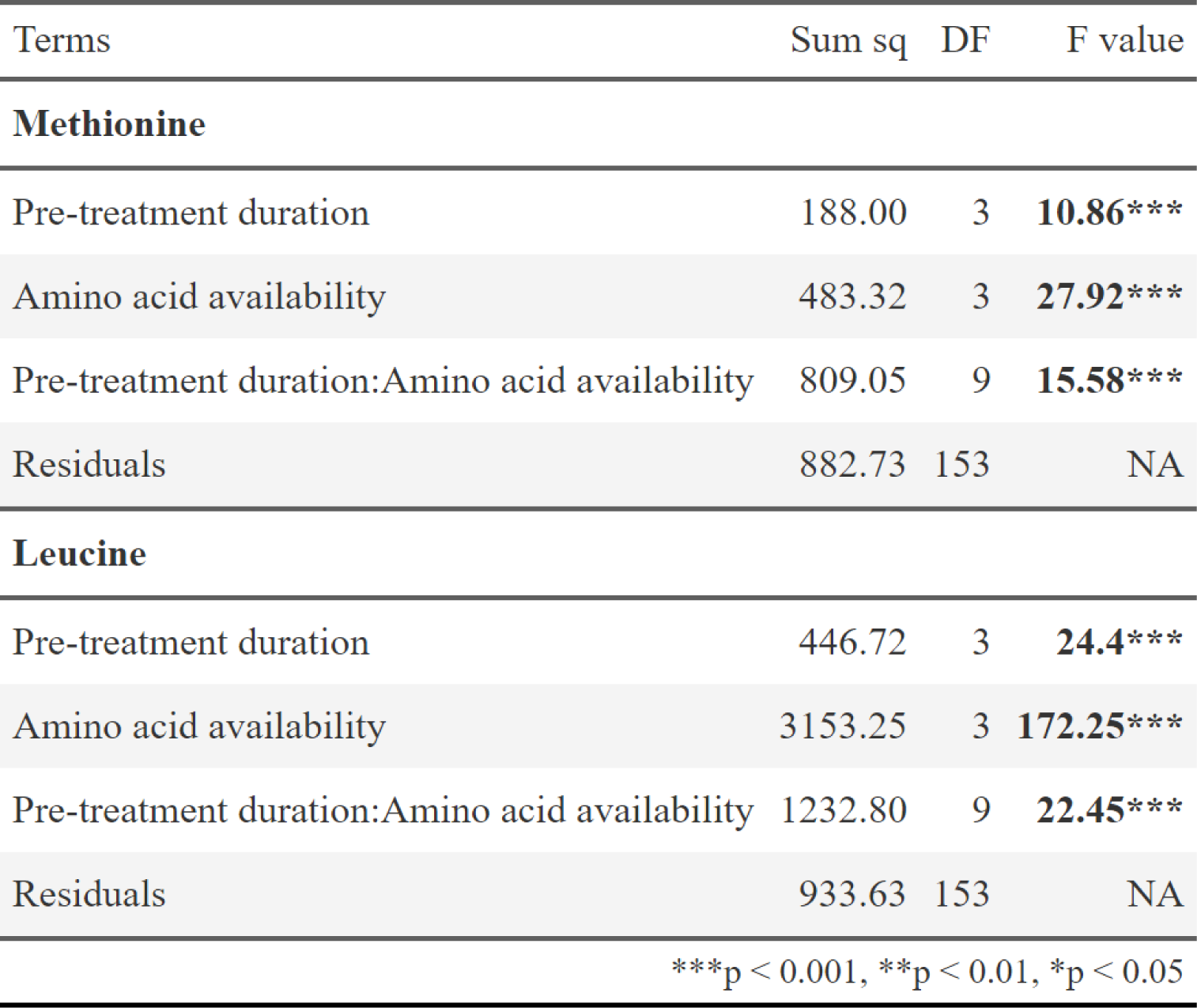
ANOVA tables for the models that best reflect the relationships between egg production, duration of pre-treatment, focal amino acid, and the level that amino acid was restricted to. Both models: Eggs laid per fly ∼ Duration * Available amino acid.

| Terms | Sum sq | DF | F value |
| --- | --- | --- | --- |
| **Methionine** |  |  |  |
| Pre-treatment duration | 188.00 | 3 | **10.86\*\*\*** |
| Amino acid availability | 483.32 | 3 | **27.92\*\*\*** |
| Pre-treatment duration:Amino acid availability | 809.05 | 9 | **15.58\*\*\*** |
| Residuals | 882.73 | 153 | NA |
| **Leucine** |  |  |  |
| Pre-treatment duration | 446.72 | 3 | **24.4\*\*\*** |
| Amino acid availability | 3153.25 | 3 | **172.25\*\*\*** |
| Pre-treatment duration:Amino acid availability | 1232.80 | 9 | **22.45\*\*\*** |
| Residuals | 933.63 | 153 | NA |
| ***$p < 0.001$, **$p < 0.01$, *$p < 0.05$ | | | |

**Table 4.** ANOVA tables for the models that best reflect the relationships between nicotine resistance, duration of pre-treatment, focal amino acid, and the level that amino acid was restricted to. Methionine model: Age ∼ poly(Duration, 2) * poly(Available methionine, 2). Leucine model: Age ∼ Duration * Available leucine.

| Terms | Sum sq | DF | F value |
| --- | --- | --- | --- |
| **Methionine** |  |  |  |
| (Intercept) | 1201158.83 | 1 | **1504.17\*\*\*** |
| poly(Pre-treatment duration, 2) | 1714.61 | 2 | 1.07 |
| poly(Available methionine, 2) | 6920.16 | 2 | **4.33\*** |
| poly(Pre-treatment duration, 2):poly(Available methionine, 2) | 11648.22 | 4 | **3.65\*\*** |
| Residuals | 671580.49 | 841 | NA |
| **Leucine** |  |  |  |
| (Intercept) | 153093.98 | 1 | **190.22\*\*\*** |
| Pre-treatment duration | 237.15 | 1 | 0.29 |
| Available leucine | 4319.50 | 1 | **5.37\*** |
| Pre-treatment duration:Available leucine | 2283.02 | 1 | 2.84 |
| Residuals | 680894.02 | 846 | NA |
| ***p < 0.001, **p < 0.01, *p < 0.05 |  |  |  |

When pre-treated flies were exposed to nicotine, there was an interaction (2^nd^ order polynomial) between the amount of methionine in the diet and how long the diet was fed to the flies that modified their survival (Table 4; Figure 4. C). The flies that lived the longest when exposed to nicotine were fed a diet of 25% methionine for 7 days before they were poisoned (mean survival 60.7h on nicotine) which was protective when compared to the complete medium (mean survival of 46.1h; P = 0.012). This result further uncouples nicotine resistance and reproductive capacity, as the fecundity of these flies did not differ from that of the fully fed controls. When leucine was modified in the diet, the duration of pre-treatment did not significantly impact survival, only the amount of leucine (Table 4, Figure 4. D). The flies that survived the longest were fed a 50% leucine diet for 5 days (mean survival 57h on nicotine), but this was not protective when compared to the complete diet (mean survival of 46.1h; P = 0.09). These results again signal that fly physiology responds in a specific way to each amino acid and the degree to which it varies, and that there is not a simple trade-off between reproduction and survival under toxic conditions.

### Diets change how flies respond to different forms of stress

Previously, we have explored how nicotine resistance can be modified by diet, though there are many ways to stress a fly, some of which have been linked to nutrition (Buchon, Silverman, and Cherry 2014; Rion and Kawecki 2007; Sgrò, Terblanche, and Hoffmann 2016; Ristow and Schmeisser 2011). Given that there is not necessarily an obvious relationship between diet and nicotine resistance, we wondered whether the diets that protect against nicotine could protect against a broad spectrum of stressors. To explore this, we selected a panel of diets that were the most protective for a focal amino acid against nicotine and used them as pre-treatment before exposure to various stressors. This panel included 7-day treatment with either an isoleucine dropout, a 25% threonine diet, or a 25% methionine diet and, although it was not protective against nicotine, we also included a 5-day, 50% leucine pre-treatment condition. As controls, we included a complete diet as a reference and a diet lacking all amino acids and only investigated the effects of these diets using female flies due to the differences we found between male and female response to diet.

#### Oxidative stress (paraquat)

Paraquat (N, N′-dimethyl-4,4′-bipyridinium dichloride) is commonly used to induce oxidative stress in *Drosophila*. Paraquat reacts in vivo to ultimately produce a superoxide anion, which is a reactive oxygen species (ROS) that can cause damage to lipids, proteins, and DNA (Suntres 2002). ROS can also be produced endogenously as a result of normal mitochondrial function (Sarniak et al. 2016), meaning that it is important for organisms to have systems to mitigate the effects of ROS. To see if we could potentially prime these systems with diet, flies were pre-treated with our panel of diets and then chronically exposed to 10mM paraquat in their food. We found that diet modified paraquat resistance (χ^2^_5_ = 60.0, P < 0.001), but only the isoleucine dropout pre-treatment protected flies against paraquat (P < 0.001; Figure 5. A; Table 5). We also found that a diet lacking all amino acids reduced flies’ capacity to resolve oxidative stress (P = 0.003). These data hint that there might be effects of isoleucine deprivation that are not elicited by other types of amino acid restriction.

**Figure 5.**
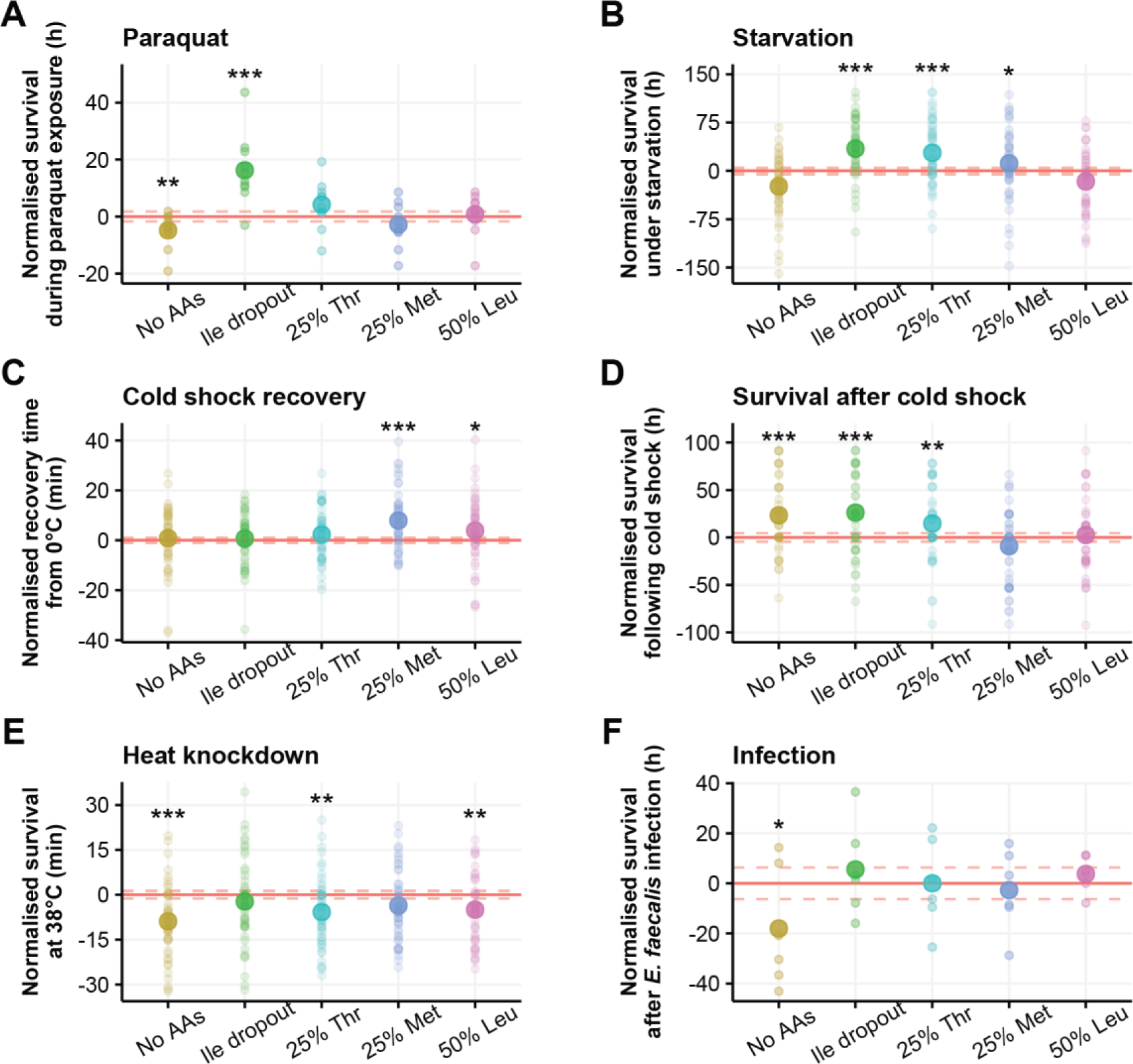
Manipulating individual dietary amino acids can differentially change how flies respond to stress. Flies were pre-treated with one of six diets before being exposed to a physical stress. Survival is represented as the difference between pre-treated flies and the controls, which were fed a nutritionally complete diet (red horizontal line +/− SE indicated by dashed red lines), small circles represent mean lifespan for each replicate and large circles represent the group mean. (**A**) Flies that were pre-treated with an isoleucine dropout were more resistant to 10mM paraquat (P < 0.001), and removing all amino acids reduced paraquat resistance (P = 0.003). (**B**) Starvation resistance was increased by pre-treating flies with an isoleucine dropout (P < 0.001) or a diet containing either 25% threonine (P < 0.001) or methionine (P < 0.05). (**C**) When cold shocked flies were transferred to room temperature, the flies that were pre-treated with either a 25% methionine diet (P < 0.001) or a 50% leucine diet (P = 0.01) took longer to recover, (**D**) However, the survival of these flies following cold shock was no different from the complete diet (P > 0.2). Survival after cold shock was improved when flies were pre-treated with a diet lacking all amino acids (P < 0.001), an isoleucine dropout diet (P < 0.001), or a 25% threonine diet (P = 0.004). (**E**) Pre-treatment diets that lack all amino acids (P < 0.001) - or contain only 25% threonine (P = 0.002) or 50% leucine (P = 0.002) - increase susceptibility to heat knockdown. (**F**) Flies that were pre-treated with a diet lacking amino acids were more susceptible to infection with *E. faecalis*. The number of individuals varied between 29-50 flies per pre-treatment group for each experiment (Table 5). ***P < 0.001, ** P < 0.01, *P < 0.05.

**Table 5.** Summary table for survival of pre-treated flies compared to control flies across multiple stressors. Control flies were fed a nutritionally complete diet. Summary of cox-proportional hazards modelling. Confidence level = 95%.

| Terms | Paraquat |  |  | Starvation |  |  | Heat knockdown |  |  | Cold recovery |  |  | Cold survival |  |  | Infection |  |  |
| --- | --- | --- | --- | --- | --- | --- | --- | --- | --- | --- | --- | --- | --- | --- | --- | --- | --- | --- |
|  | n | Estimate | Z value | n | Estimate | Z value | n | Estimate | Z value | n | Estimate | Z value | n | Estimate | Z value | n | Estimate | Z value |
| Complete diet | 48 | NA | NA | 41 | NA | NA | 49 | NA | NA | 48 | NA | NA | 50 | NA | NA | 30 | NA | NA |
| No AAs | 48 | 0.601 | **2.95\*\*** | 44 | 0.349 | 1.58 | 49 | 1.137 | **5.46\*\*\*** | 49 | -0.108 | -0.53 | 50 | -1.155 | **-3.5\*\*\*** | 30 | 0.845 | **2.14\*** |
| Ile dropout | 49 | -0.936 | **-4.4\*\*\*** | 48 | -1.230 | **-5.14\*\*\*** | 49 | 0.182 | 0.89 | 47 | -0.060 | -0.29 | 50 | -1.529 | **-4.04\*\*\*** | 30 | -0.292 | -0.62 |
| 25% Thr | 50 | -0.357 | -1.76 | 48 | -0.861 | **-3.84\*\*\*** | 50 | 0.637 | **3.14\*\*** | 47 | -0.359 | -1.74 | 49 | -0.887 | **-2.83\*\*** | 30 | 0.074 | 0.17 |
| 25% Met | 50 | 0.273 | 1.35 | 48 | -0.559 | **-2.54\*** | 49 | 0.313 | 1.53 | 48 | -0.920 | **-4.37\*\*\*** | 50 | 0.315 | 1.3 | 29 | 0.120 | 0.28 |
| 50% Leu | 47 | -0.039 | -0.19 | 46 | 0.284 | 1.31 | 50 | 0.628 | **3.08\*\*** | 50 | -0.520 | **-2.56\*** | 50 | -0.044 | -0.18 | 30 | -0.075 | -0.17 |

\*\*\*p < 0.001, \*\*p < 0.01, \*p < 0.05

#### Starvation

A well-established method for increasing starvation resistance in flies is nutrient restriction. This could be in the form of dietary restriction (Chippindale, Chu, and Rose 1996), protein restriction (Leroi, Kim, and Rose 1994), or even single amino acid deprivation (Srivastava et al. 2022). This is hypothesised to result from flies responding to these diets by reducing their reproductive output and in turn, storing nutrients such as fats which can be used to maintain life when starved (Rion and Kawecki 2007). When we starved flies after our single amino acid pre-treatments, we found that diet affected survival (χ^2^_5_ = 74.9, P < 0.001). Specifically, we found that an isoleucine dropout diet, a 25% threonine diet, or a 25% methionine diet increased starvation resistance (Figure 5. B; Table 5). Flies that were pre-treated with either a protein free diet or a 50% leucine diet responded to starvation no differently than the fully fed control flies. Given that we have shown that a 25% methionine diet does not reduce fecundity and increases starvation resistance, and that a diet lacking amino acids reduces fecundity but does not protect against starvation, our data suggest that fecundity is not simply traded for starvation resistance.

#### Cold shock

In nature, organisms are faced with thermal challenges such as extremely cold or hot conditions to which they must evolve resistance or tolerance. Time to wake up after chill coma is a trait that is often measured in the context of population plasticity in the face of climate change (David et al. 1998), but subsequent survival in the following days is not always measured. Here, we measured both. We found that pre-treatment diet impacted both recovery time (χ^2^_5_ = 28.8, P < 0.001) and survival following cold shock (χ^2^_5_ = 53.2, P < 0.001). However, none of our pre-treatment diets improved chill-coma recovery time (Figure 5. C; Table 5), in fact, a 25% methionine or a 50% leucine diet increased the time it took for flies to wake. However, we found that flies pre-treated with an isoleucine dropout, a 25% threonine, or a protein-free diet lived longer in the 5 days following cold shock (Figure 5. D; Table 5). When we examined the data across all pre-treatment groups, we discovered that there is an overall correlation between recovery time and survival post cold shock (P < 0.001, Pearson’s correlation coefficient = −0.24). However, the pre-treatment diets that improved survival following cold shock did not differ in their recovery time from the complete diet. These results highlight that flies can recover at the same rate from chill coma, but this is not necessarily an indication of their health status, as many die in the following days. They also indicate that pre-treatment diets can improve the health status of flies following cold shock without influencing recovery time.

#### Heat knockdown

On the other end of the thermal spectrum from cold shock is heat shock. In the face of climate change, animals are facing increasing temperatures and so understanding their upper thermal limit is an important indicator of stress resistance with implications for population persistence (Hoffmann, Chown, and Clusella-Trullas 2013). It is more typical however to manipulate the diet of flies during their development than during adulthood, and there is a link between larval diet and upper thermal tolerance (Andersen et al. 2010; Sisodia and Singh 2012). We were therefore interested to know whether there is a similar link between heat tolerance and our adult pre-treatment diets. We found that pre-treatment diet affected heat knockdown time (χ^2^_5_ = 35.3, P < 0.001), though none of our pre-treatment diets enhanced heat tolerance when compared to the complete diet (Figure 5. E; Table 5). In fact, flies that were pre-treated with a 25% threonine diet or a 50% leucine diet were more susceptible to heat knockdown than the complete diet. These results indicate that the mechanisms by which these diets are protecting flies against nicotine are not generalisable across all types of stress.

#### Infection with E. faecalis

We were also interested to know whether our diets could protect flies against a biotic stress. To do this, we opted to prick flies with live *Enterococcus faecalis*, a bacterium that naturally colonises the gastrointestinal tract of flies (Cox and Gilmore 2007). Enterococci are a leading cause of nosocomial infections in humans (Sood et al. 2008), and *E. faecalis* is routinely used to infect *Drosophila* and study host-pathogen interactions (Lazzaro, Sackton, and Clark 2006; Chapman et al. 2020; Cabrera et al. 2023). When we infected flies with *E. faecalis*, we found that pre-treatment diet overall did not impact survival (χ^2^_5_ = 9.6, P < 0.09). Though when we compared survival between different pre-treatment diets and the complete diet, we found that a diet lacking amino acids increased susceptibility to the pathogen (Figure 5. F; Table 5). When we infected the flies, we were also interested to know whether pre-treatment diet could impact fecundity under infection, as we expected that the energy that would be expended on laying eggs would need to be diverted to resolve the infection. However, we did not see any evidence for this, and flies that were infected had indistinguishable egg laying dynamics to control flies for each pre-treatment (Figure 6; Table 6; Supplementary table 5). Together, these results show that the ways that diet can protect against nicotine poisoning are not the same for infection with *E. faecalis*.

**Figure 6.**
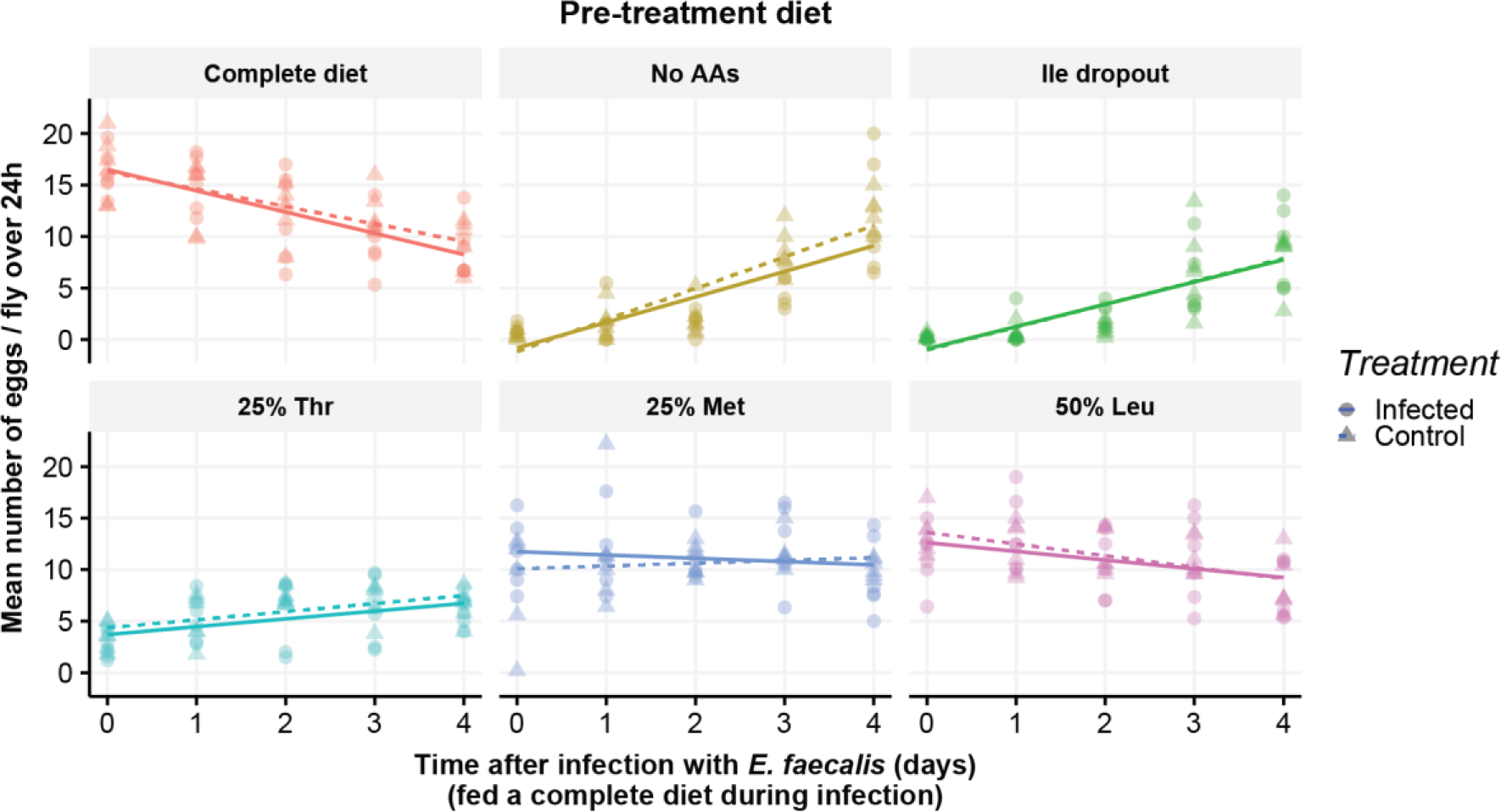
Pre-treatment diet did not influence fecundity during infection. Flies were pre-treated with one of six diets before being infected with *E. faecalis* and transferred onto a complete diet. The trends in egg laying between infected (solid lines) and control (dashed lines) flies were not different from each other across the pre-treatment diets. The number of individuals for each condition was 30 flies.

**Table 6.** ANOVA table for the model that looks for interactions between the number of eggs laid by infected and control females: Eggs laid per fly ∼ Pre-treatment diet * Infection status * Time after infection.

| Terms | Sum sq | DF | F value |
| --- | --- | --- | --- |
| (Intercept) | 2723.49 | 1 | **348.51\*\*\*** |
| Pre-treatment diet | 2786.36 | 5 | **71.31\*\*\*** |
| Infection status | 0.21 | 1 | 0.03 |
| Time after infection | 255.81 | 1 | **32.73\*\*\*** |
| Pre-treatment diet:Infection status | 21.21 | 5 | 0.54 |
| Pre-treatment diet:Time after infection | 942.11 | 5 | **24.11\*\*\*** |
| Infection status:Time after infection | 4.14 | 1 | 0.53 |
| Pre-treatment diet:Infection status:Time after infection | 18.07 | 5 | 0.46 |
| Residuals | 2625.76 | 336 | NA |

\*\*\*p < 0.001, \*\*p < 0.01, \*p < 0.05

## Discussion

In this manuscript, we showed that there is sexual dimorphism in the protection afforded by short-term individual amino acid deprivation, and that the protection cannot be mimicked by manipulating the total protein in the diet. Moreover, we found further evidence that there are different optimal pre-treatments for each amino acid to increase stress resistance of the consumer. Finally, we showed that different pre-treatment diets could protect flies against different stressors to different extents. This research furthers our understanding of the benefits that are conferred by amino acid-restricted diets and offers insights into the complex relationship between nutrition and stress resistance.

In our previous work, we found that diets lacking an essential amino acid protected female flies from subsequent nicotine poisoning. It is particularly common in studies that investigate diet using fruit flies to only experiment with females, as changes in diet lead to rapid, observable changes in fecundity (Sang and King 1961). Several studies have also observed larger effect sizes of diet on females than males which is generally attributed to females eating more than males to sustain their reproductive output (Partridge, Piper, and Mair 2005; Wong et al. 2009). Given this information, we assumed that males would also receive the benefits of an isoleucine dropout, but perhaps to a lesser degree than females. This was not the case however, and instead we found that short-term isoleucine deprivation reduced the nicotine resistance of males. Interestingly, fully-fed males were more tolerant to nicotine than fully-fed females, which is surprising given that young females typically have greater resistance to a range of stressors than the same age males (Belyi et al. 2020). This could reflect a difference in initial investment capabilities of the sexes, as the females in our experiments were mated and so were committed to a greater reproductive investment than the males.

It is also possible that these differences could reflect the difference in food, and so toxin, consumption by male and female flies (Lee, Kim, and Min 2013). Differentiating between these possibilities could be addressed by administering nicotine using capillary feeders that would permit precise quantification of toxin ingestion (Ja et al. 2007; Diegelmann et al. 2017). It is also possible that the combinations that benefit females simply do not benefit males, and that the sexes have different responses to diet. Given that the identity of the focal amino acids, as well as the degree of restriction and pre-treatment duration all interact to impact female nicotine resistance, it is likely that sex also modifies this response.

To understand this, we should investigate the differences in molecular responses to isoleucine deprivation between males and females. Previous research has shown sex-specific transcriptional responses across high and low protein diets, implying that the output of nutrient sensing pathways is different between male and female flies (Camus, Piper, and Reuter 2019). It is important that we understand the physiological differences in responses to diet between sexes if we are to make any suggestions about diet improving human health in the future.

The benefits of chronic protein restriction, including both lifespan and health span extension, have been documented across a wide range of taxa (Mirzaei, Suarez, and Longo 2014). Recently, the acute benefits of protein restriction have also come to light. For example, mice are more resistant to hepatic and renal ischaemic reperfusion injury, which are models of liver and kidney surgery respectively, when they have been fed protein restricted pre-treatment diets (Harputlugil et al. 2014; Robertson et al. 2015). Similarly, protein restriction protects flies against hydrogen peroxide, an oxidising agent, and also protects old flies against infection with *E. faecalis* (Zhang et al. 2023). Since restricting an individual amino acid protects flies against nicotine, it was surprising to find that protein restriction by restricting all dietary amino acids did not also protect flies. It is possible, therefore, that flies require one or more of the remaining non-focal amino acids to be supplied in the diet to establish resistance.

A potential contender for this amino acid is leucine, since no methods of leucine restriction protected flies against nicotine. It would be interesting to know if restricting all amino acids except leucine is beneficial, or if the benefits of combining nicotine-protective diets (ie. food lacking isoleucine, with 25% threonine, 25% methionine, and containing leucine) is more protective than any single amino acid restriction alone. The current framework of research mainly focuses on observing the protective effects of protein or dietary restriction (Emran et al. 2014; Joußen et al. 2008), but our work emphasises that dietary restriction and individual amino acid restriction are not equivalent, and sometimes amino acid restriction is protective when other types of restriction are not. Thus, there is merit in altering the levels of individual dietary amino acids even when there is no, or negative, effects of other dietary restrictions.

A common theme from our data is that short term isoleucine deprivation was the most common way to protect flies against stressors. Isoleucine restriction has recently garnered more attention than that of other amino acids, because of the generally protective effects that result from restricting its intake. When it is restricted in the diet, female flies live longer (Weaver et al. 2023) and so do male and female heterogenous mice, who also have improved metabolic health (Green et al. 2023). We also recently found that female flies subjected to two short bouts of isoleucine deprivation have extended lifespan (Fulton et al. 2024). A potential explanation for this is that isoleucine deprivation triggers protective systems that bolster defence against a broad spectrum of damage threats and that these mechanisms overlap with those that improve lifespan. One possibility is a link between detoxification capacity and longevity; detoxification genes are strongly upregulated in long-lived insulin mutants (McElwee et al. 2007) and insulin mutant flies are more resistant to DDT (Gronke et al. 2010). Since flies were not protected by removing isoleucine at the same time as the other amino acids (ie. no protein treatment), we propose that protection requires new protein synthesis. New proteins would surely require isoleucine and this could possibly be made available by recycling amino acid stores via protein breakdown (Johnstone et al. 2024). If this is the case, then these stored amino acids must somehow be reserved for somatic protection, rather than for use in egg production, which ceases when any essential amino acid is removed from the diet (Sang and King 1961; Alves et al. 2022). Studies tracking the fate of labelled amino acids into protein during isoleucine deprivation could be revealing, both for understanding the systems that are triggered to enhance stress resistance as well as to identify protein synthesis that is required to sustain lifespan.

Another factor that animals encounter in the wild is changes in temperature. It is particularly important for ectotherms to be aware of cues that signify a temperature change since they cannot regulate their own body temperature, and so must respond physiologically to ensure survival (Angilletta Jr, Niewiarowski, and Navas 2002). Interestingly, we found nicotine-protective diets that reduced fecundity also increased survival following cold shock. This trade-off could be explained by the availability of sterols, essential micronutrients that modulate cell membrane fluidity (Dufourc 2008), which are speculated to be important in mitigating mechanical injury to cell membranes during cold shock (Teets and Denlinger 2013). Flies that are fed more cholesterol during development can have enhanced cold tolerance (Shreve, Yi, and Lee 2007; Allen et al. 2024) and the trade-off between lifespan and reproduction can be rescued by supplementing cholesterol (Zanco et al. 2021), allowing flies to have long lives and high fecundity. Taken together, it is possible that reduced fecundity due to essential amino acid deprivation increases body sterol stores, and this modifies cell membranes in a way that protects flies against acute cold shock. If this were the case, we would expect that supplementing adult flies with cholesterol would also protect them against cold shock, even when fed a high protein diet.

We also found that there was some overlap between diets that protected against survival following a cold shock and starvation. This could be explained by the changing of seasons. Winter is associated with reduced temperatures and availability of nutrition and evolving resistance to both simultaneously should be adaptive. Interestingly, flies that are evolved for chill coma recovery have higher levels of phosphatidic acids, but not triacylglyceride (TAG) or lipid levels (Ko et al. 2019). We found that the TAG levels of flies that were pre-treated with an isoleucine dropout or a 25% threonine diet were not different from controls fed a complete diet, but both of these pre-treatment diets protect against starvation and cold stress. It would be interesting to look at the levels of phosphatidic acids in our pre-treated flies as a possible mechanism for cross-protection against starvation and cold stress.

Previously, we have found that rapamycin, a drug that inhibits Target of Rapamycin (TOR), protected fully-fed flies against nicotine to the same degree as isoleucine deprivation (Fulton, Mirth, and Piper 2022). As TOR is a master regulator of growth, which detects cellular levels of amino acids and is inactivated by low levels of amino acids (Saxton and Sabatini 2017), we assumed that both isoleucine deprivation and rapamycin achieved nicotine resistance through TOR suppression. However, other studies have shown that rapamycin treatment improves heat tolerance of wildtype flies (Willot et al. 2023) and can improve the maximum thermal temperature (CTmax) withstood by DGRP flies (Rohde et al. 2021). We therefore anticipated that short term isoleucine deprivation would also increase heat tolerance in our flies, but it did not. Interestingly, other work has also shown that flies under dietary protein restriction are less heat tolerant (Emran et al. 2014). While dietary protein restriction and rapamycin treatment are thought to increase lifespan through TOR suppression (Partridge et al. 2011; Kapahi, Kaeberlein, and Hansen 2017), these data suggest that the molecular changes induced by rapamycin treatment are overlapping with those induced by isoleucine or protein restriction, but not the same. It would be interesting to investigate the molecular responses to isoleucine deprivation and rapamycin treatment to understand how their overlaps and differences shape the way these treatments enhance resistance to various stressors as well as enhance lifespan.

## Conclusion

Our study sheds light on the intricate relationship and trade-offs between nutrition and stress resistance in *Drosophila melanogaster*. We have demonstrated that short-term amino acid restrictions, particularly isoleucine deprivation, can protect against various stressors, including nicotine poisoning, oxidative stress, starvation, and cold shock. We observed sexual dimorphism in the response to dietary manipulation, with females, but not males, benefiting from isoleucine deprivation. Our findings also highlight the differences between total protein restriction and individual amino acid restriction, emphasising the importance of considering specific amino acids in diet manipulation studies. Future investigations into the molecular responses to dietary interventions and their implications for stress resistance across multiple stressors will provide valuable insights into optimising health span and resilience across species.

## Acknowledgements

We would like to thank Professor Damian Dowling and Professor Carla Sgro for sharing their equipment for heat and cold shocking flies, and Dr Winston Yee for assisting us to set up these thermal tolerance assays.

## Supplementary materials

**Supplementary table 1.** Ingredients in 1L of Sugar Yeast (SY) medium.

| Ingredient | Amount (L<sup>-1</sup>) | Supplier (catalogue number) |
| --- | --- | --- |
| Sugar | 50g | Bundaberg Australia (M180919) |
| Autolysed Brewer's yeast | 100g | MP Biomedicals (290331225) |
| Agar (grade J3) | 10g | Gelita Australia (A-181017) |
| Nipagin (10% w/v in 96% EtOH) | 30mL | Sigma Aldrich (W271004) |
| Propionic acid | 3mL | Merck (8.00605) |

**Supplementary table 2.** Ingredients in 1L of complete synthetic medium.

| Ingredient | Amount (L<sup>-1</sup>) | Supplier (catalogue number) |
| --- | --- | --- |
| L-arginine HCl | 0.814g | Sigma Aldrich (A5131) |
| L-alanine | 0.551g | Sigma Aldrich (A7627) |
| L-asparagine | 0.514g | Sigma Aldrich (A0884) |
| L-aspartic acid | 0.586g | Sigma Aldrich (A6683) |
| L-cysteine | 0.171g | Sigma Aldrich (C7477) |
| L-glutamic acid | 0.759g | Sigma Aldrich (G5889) |
| L-glutamine | 0.560g | Sigma Aldrich (G3126) |
| Glycine | 0.383g | Sigma Aldrich (G7126) |
| L-histidine | 0.327g | Sigma Aldrich (H8000) |
| L-isoleucine | 0.560g | Sigma Aldrich (I2752) |
| L-leucine | 1.020g | Sigma Aldrich (L8912) |
| L-lysine HCl | 0.682g | Sigma Aldrich (L5626) |
| L-methionine | 0.301g | Sigma Aldrich (M9625) |
| L-phenylalanine | 0.504g | Sigma Aldrich (P2126) |
| L-proline | 0.489g | Sigma Aldrich (P0380) |
| L-serine | 0.688g | Sigma Aldrich (S4500) |
| L-threonine | 0.552g | Sigma Aldrich (T8625) |
| L-tryptophan | 0.160g | Sigma Aldrich (T0254) |
| L-tyrosine | 0.460g | Sigma Aldrich (T8566) |
| L-valine | 0.599g | Sigma Aldrich (V0500) |
| Agar | 7.00g | Sigma Aldrich (A7002) |
| Sucrose | 17.12g | Sigma Aldrich (S1888) |
| Cholesterol | 0.3g | Glentham Life Sciences (GE0100) |
| Choline chloride | 0.05g | Sigma Aldrich (C1879) |
| Myo-inositol | 0.005g | Sigma Aldrich (I7508) |
| Inosine | 0.065g | Sigma Aldrich (I4125) |
| Uridine | 0.060g | Sigma Aldrich (U3750) |
| Thiamine | 0.0014g | Sigma Aldrich (T4625) |
| Riboflavin | 0.0007g | Sigma Aldrich (R4500) |
| Nicotinic acid | 0.0084g | Sigma Aldrich (N4126) |
| Ca pantothenate | 0.0108g | Sigma Aldrich (21210) |
| Pyridoxine-HCl | 0.0017g | Sigma Aldrich (P9755) |
| Biotin | 0.0001g | Sigma Aldrich (B4501) |
| Folic acid | 0.0005g | Sigma Aldrich (F7876) |
| CaCl<sub>2</sub>.2H<sub>2</sub>O | 0.250g | Sigma Aldrich (C7902) |
| CuSO<sub>4</sub>.5H<sub>2</sub>O | 0.0025g | Sigma Aldrich (C7631) |
| FeSO<sub>4</sub>.7H<sub>2</sub>O | 0.025g | Sigma Aldrich (F7002) |
| MgSO<sub>4</sub> (anhydrous) | 0.250g | Sigma Aldrich (M7506) |
| MnCl<sub>2</sub>.4H<sub>2</sub>O | 0.001g | Sigma Aldrich (M3634) |
| Zn SO<sub>4</sub>.7H<sub>2</sub>O | 0.025g | Sigma Aldrich (Z0251) |
| KH<sub>2</sub>PO<sub>4</sub> | 3.00g | Sigma Aldrich (P9791) |
| NaHCO<sub>3</sub> | 1.00g | Sigma Aldrich (S8875) |
| Acetic acid (glacial) | 3mL | Merck KGaA (100063) |
| Propionic acid | 6mL | Merck KGaA (8.00605) |
| Nipagin (10% w/v in 96% EtOH) | 15mL | Sigma Aldrich (W271004) |

**Supplementary table 3.**
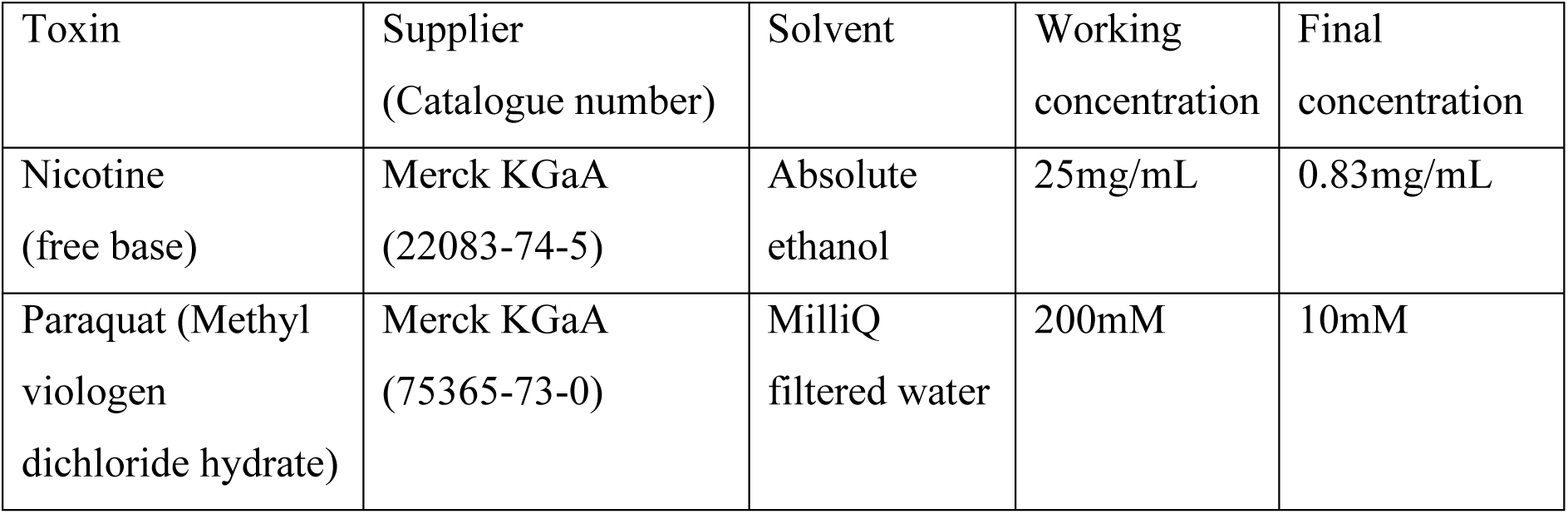
Details for the toxins used in experiments. Nicotine laced vials were prepared by aliquoting 100µL of diluted nicotine (in absolute ethanol, 25mg/mL) onto 3mL of cooled, complete synthetic food. Paraquat laced media were prepared by aliquoting 100µL of diluted paraquat (in water, 200mM) onto 2mL of cooled, complete synthetic food.

**Supplementary table 4.**
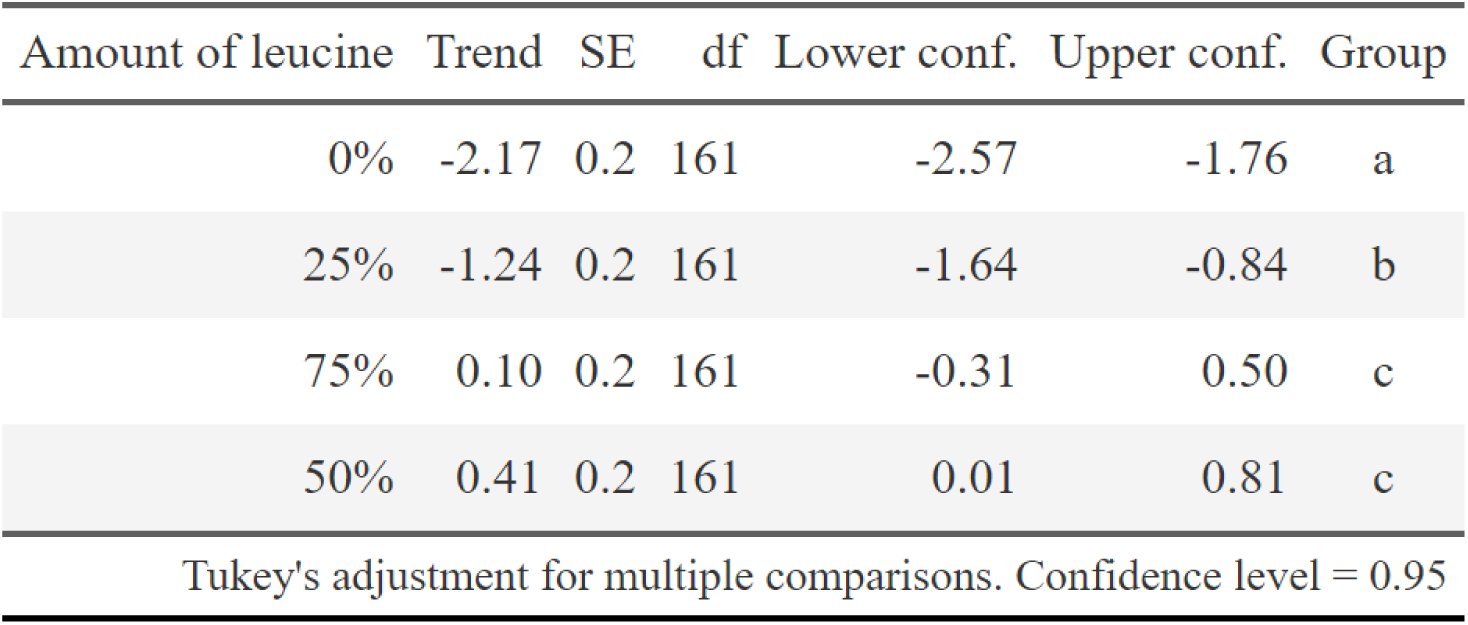
Trends in egg laying over time spent on leucine restricted diets. Estimates have been generated using emtrends, and the grouping was determined using multcomp’s compact letter display. Conditions belonging to the same group have not been shown to be the same, only that they are not different. Confidence level = 95%.

**Supplementary table 5.**
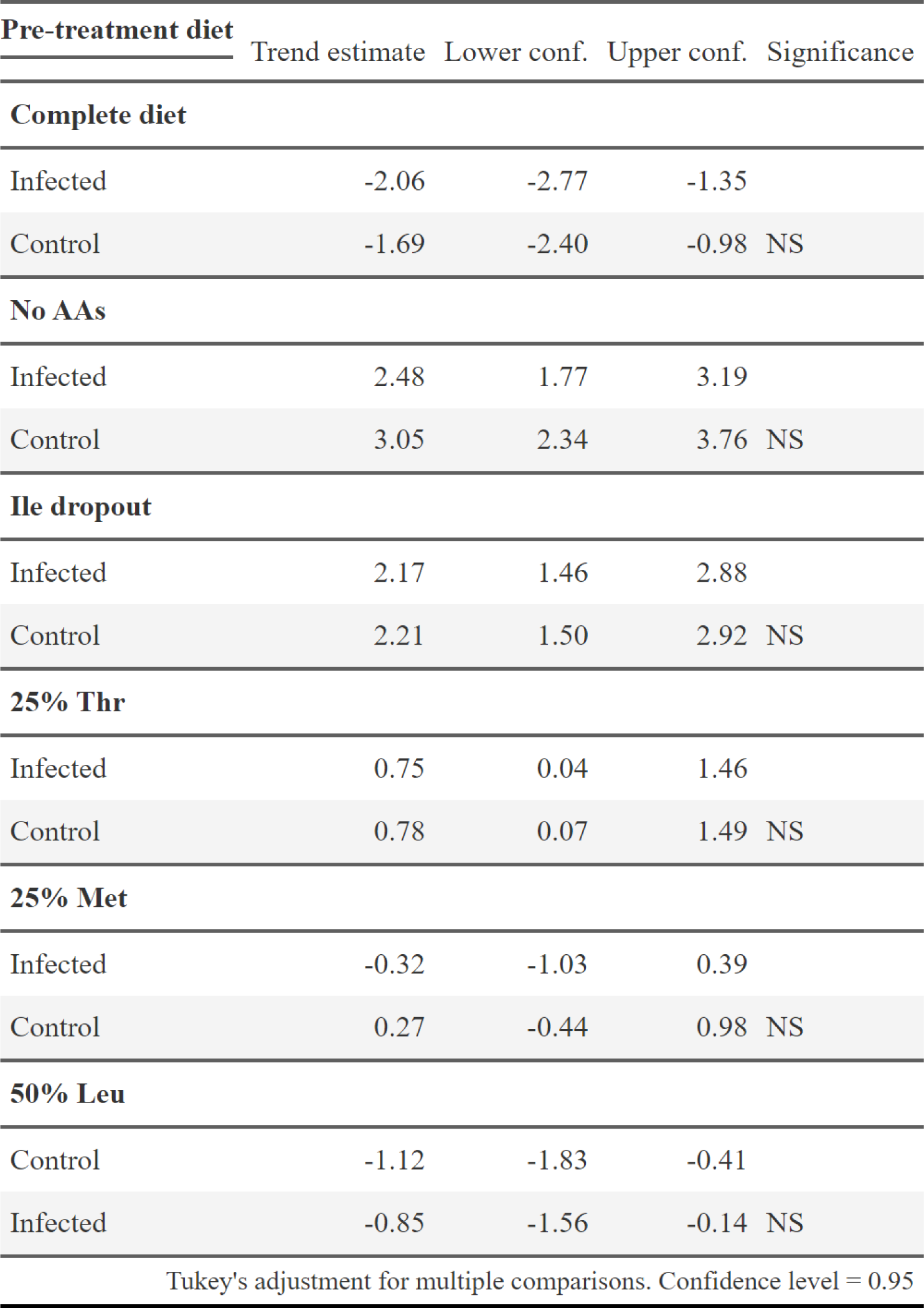
The estimated linear egg laying trend for infected and control flies, separated by pre-treatment diet. Estimates generated by emtrends (Lenth 2021). Confidence level = 95%.

| **Pre-treatment diet** | Trend estimate | Lower conf. | Upper conf. | Significance |
| --- | --- | --- | --- | --- |
| **Complete diet** |  |  |  |  |
| Infected | -2.06 | -2.77 | -1.35 |  |
| Control | -1.69 | -2.40 | -0.98 | NS |
| **No AAs** |  |  |  |  |
| Infected | 2.48 | 1.77 | 3.19 |  |
| Control | 3.05 | 2.34 | 3.76 | NS |
| **Ile dropout** |  |  |  |  |
| Infected | 2.17 | 1.46 | 2.88 |  |
| Control | 2.21 | 1.50 | 2.92 | NS |
| **25% Thr** |  |  |  |  |
| Infected | 0.75 | 0.04 | 1.46 |  |
| Control | 0.78 | 0.07 | 1.49 | NS |
| **25% Met** |  |  |  |  |
| Infected | -0.32 | -1.03 | 0.39 |  |
| Control | 0.27 | -0.44 | 0.98 | NS |
| **50% Leu** |  |  |  |  |
| Control | -1.12 | -1.83 | -0.41 |  |
| Infected | -0.85 | -1.56 | -0.14 | NS |
| Tukey's adjustment for multiple comparisons. Confidence level = 0.95 |  |  |  |  |

**Supplementary Figure 1.**
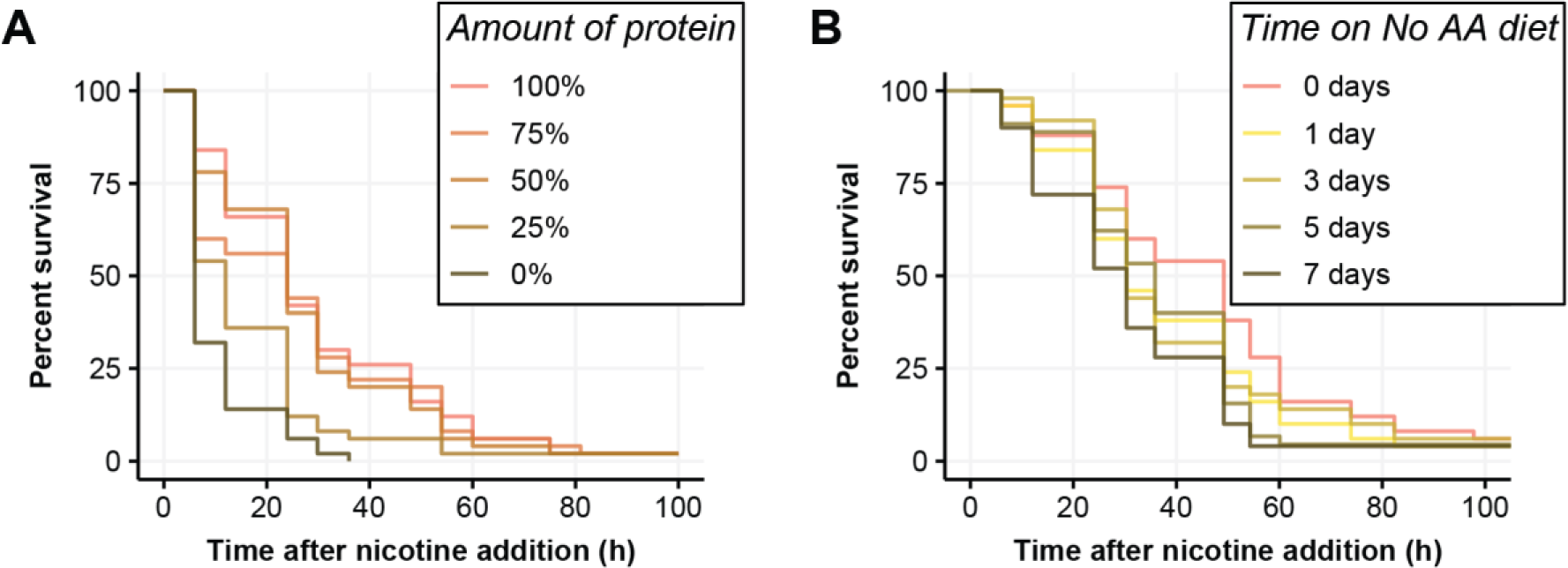
Flies were either pre-treated with diets where all amino acids were reduced for 7 days, or where all amino acids were absent for 1, 3, 5 or 7 days before chronic exposure to 0.83mg/mL nicotine. (**A**) Survival curves of flies when pre-treated with diets that have altered amino acid concentrations, survival was reduced when flies were pre-treated with 25% or 0% amino acids (P < 0.001). (**B**) Survival curves when flies were pre-treated with no amino acids for 0, 1, 3, 5 or 7 days before nicotine exposure. Flies were more susceptible to nicotine when fed no amino acids for 7 days before poisoning.

**Supplementary Figure 2.**
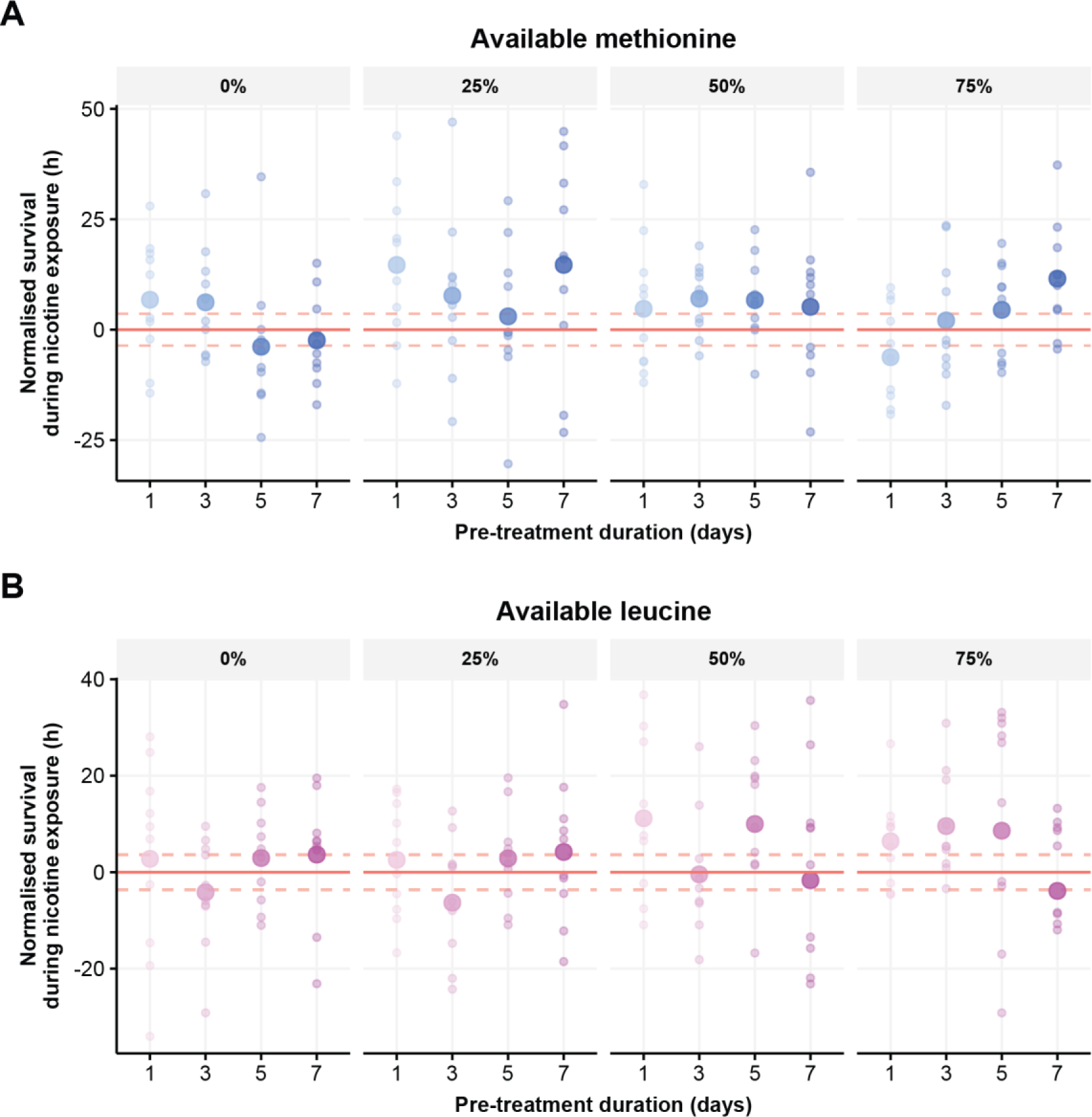
Identity of amino acid interacts with level of restriction and duration of pre-treatment to respond to nicotine poisoning. Flies were pre-treated with diets containing 1 of 5 amounts of either methionine or leucine (0%, 25%, 50%, 75%, 100%) for either 7, 5, 3, or 1 day before chronic exposure to 0.83mg/mL nicotine. (**A**) Survival time of pre-treated flies that has been normalised to survival of flies fed 100% of amino acids (red horizontal line +/− SE indicated by dashed red lines), small circles represent mean lifespan for each replicate and large circles representing the group mean. Survival when methionine was modified was influenced by both the amount of methionine and the duration of pre-treatment (Table 4), with a 25% methionine for 7 days offering the greatest protection (P = 0.012) (**B**) Leucine dose (P = 0.02) but not pre-treatment duration (P = 0.59) significantly impacted survival on nicotine, although the longest lived flies - that were fed a 50% leucine diet for 5 days - were not protected (P = 0.09). N = 50 flies per pre-treatment condition.

**Supplementary Figure 3.**
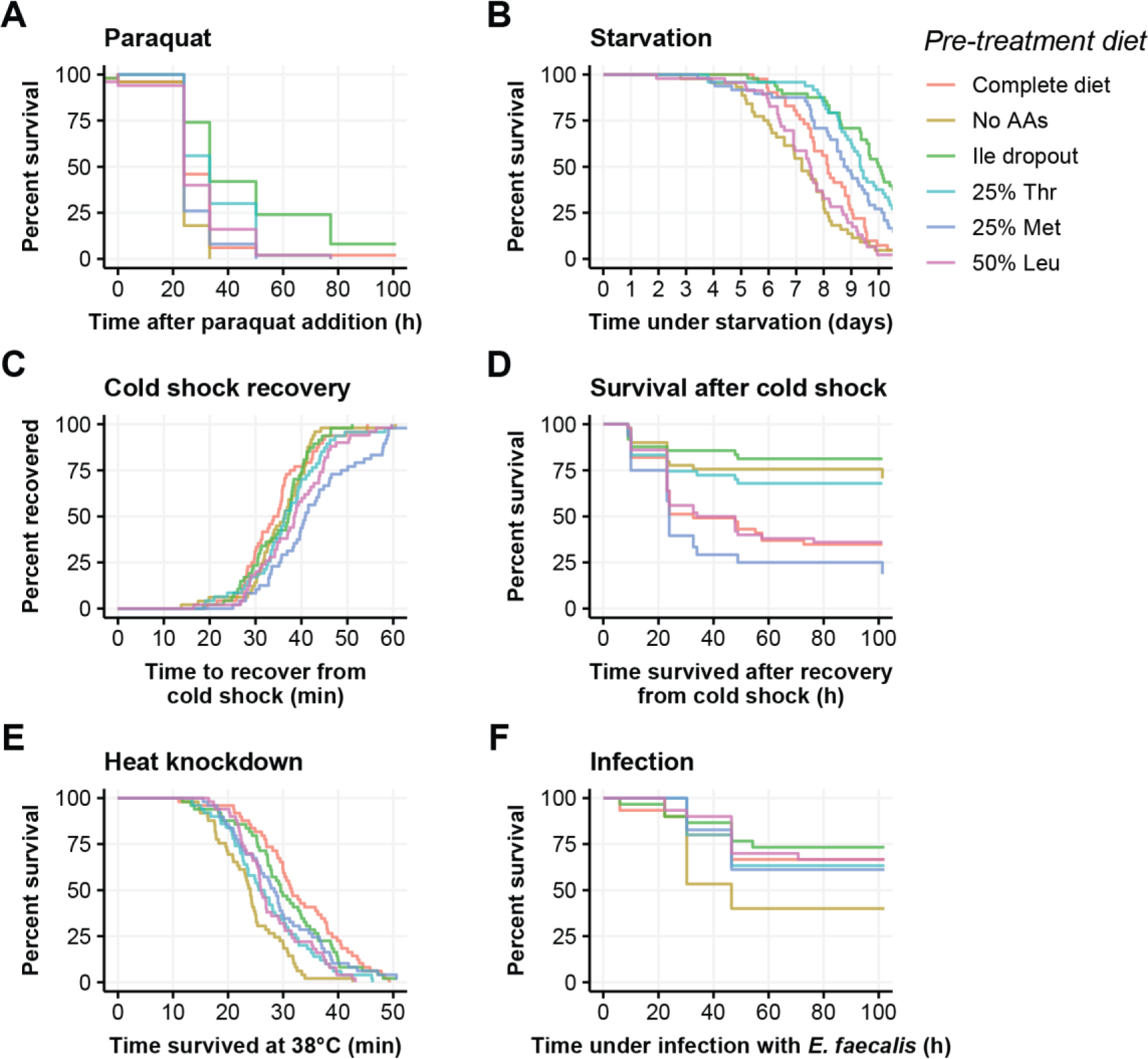
Survival curves of the data presented in Figure 5. Flies were pre-treated with one of six diets before being exposed to a physical stress. (**A**) Flies that were pre-treated with an isoleucine dropout were more resistant to 10mM paraquat (P < 0.001), and removing all amino acids reduced paraquat resistance (P = 0.003). (**B**) Starvation resistance was increased by pre-treating flies with an isoleucine dropout (P < 0.001) or a diet containing either 25% threonine (P < 0.001) or methionine (P < 0.05). (**C**) When cold shocked flies were transferred to room temperature, the flies that were pre-treated with either a 25% methionine diet (P < 0.001) or a 50% leucine diet (P = 0.01) took longer to recover, (**D**) However, the survival of these flies following cold shock was no different from the complete diet (P > 0.2). Survival after cold shock was improved when flies were pre-treated with a diet lacking all amino acids (P < 0.001), an isoleucine dropout diet (P < 0.001), or a 25% threonine diet (P = 0.004). (**E**) Pre-treatment diets that lack all amino acids (P < 0.001) - or contain only 25% threonine (P = 0.002) or 50% leucine (P = 0.002) - increase susceptibility to heat knockdown. (**F**) Flies that were pre-treated with a diet lacking amino acids were more susceptible to infection with *E. faecalis*. The number of individuals varied between 29-50 flies per pre-treatment group for each experiment (Table 5).

